# EGFR upregulation drives signaling reactivation during EGFR inhibition in glioblastoma without broad kinome rewiring

**DOI:** 10.64898/2026.08.13.744581

**Authors:** Yoran Broersma, Megan Houweling, Tsz Ting Wong, Pragallabh Purwar, Richard de Goeij de Haas, Alex A. Henneman, Sander R. Piersma, Thang V. Pham, Connie R. Jimenez, David Noske, Alan Gerber, Bart A. Westerman

## Abstract

**Background:** Epidermal growth factor receptor (EGFR) amplification occurs in ∼50% of IDH-wildtype glioblastoma (GBM) cases, frequently accompanied by expression of the oncogenic EGFRvIII variant. Although EGFR represents an attractive therapeutic target, EGFR-directed therapies have shown limited clinical efficacy in GBM. Resistance to kinase inhibitors is frequently attributed to activation of compensatory signaling pathways (“kinome rewiring”). We therefore investigated whether EGFR inhibition in GBM induces broad adaptive kinase responses that could be co-targeted to overcome resistance.

**Methods:** We molecularly profiled 29 patient-derived GBM cell lines for EGFR status and selected five representative models spanning EGFR amplification states for functional analyses. Cells were treated with EGFR inhibitors and responses were assessed using viability assays, time-resolved immunoblotting, and phosphoproteomics (LC-MS/MS) with kinase activity inference.

**Results:** EGFR inhibitors preferentially impaired viability in EGFR-driven models and transiently reduced EGFR phosphorylation during the initial response. However, partial restoration of EGFR phosphorylation and downstream signaling occurred after 24 hours of inhibitor exposure. Phosphoproteomics revealed no evidence of broad kinome rewiring within this timeframe but instead identified increased EGFR abundance, associated with partial restoration of EGFR pathway activity. The phosphorylated-to-total EGFR ratio remained stable, indicating that increased EGFR abundance may enable persistent residual kinase activity despite continued, but incomplete, target inhibition.

**Conclusions:** Early responses to EGFR inhibition in GBM were not characterized by broad kinome rewiring but by restoration of EGFR signaling associated with increased EGFR abundance. These findings suggest that adaptive signaling remains largely EGFR-dependent despite inhibitor exposure, identifying regulation of EGFR abundance as a potential contributor to therapeutic resistance.

**Key points:**

- Early responses to EGFR inhibition occur without evidence of broad kinome rewiring.
- EGFR signaling is restored during sustained inhibitor exposure.
- Increased EGFR abundance is associated with restoration of pathway activity.

**Importance of the study:** Adaptive resistance to EGFR-targeted therapies in GBM is commonly attributed to activation of alternative signaling pathways. Using patient-derived GBM models and phosphoproteomic profiling, we show that early adaptive responses to EGFR inhibition are not characterized by broad kinome signaling rewiring but instead remain centered on reactivation of EGFR signaling. Our findings suggest that increased EGFR abundance in response to inhibitor exposure may enhance residual EGFR signaling sufficiently to partially restore downstream pathway activity. These results indicate that early adaptive responses to EGFR inhibition may remain largely EGFR-dependent, potentially limiting the effectiveness of strategies primarily aimed at co-targeting alternative signaling pathways.

**Graphical abstract:** 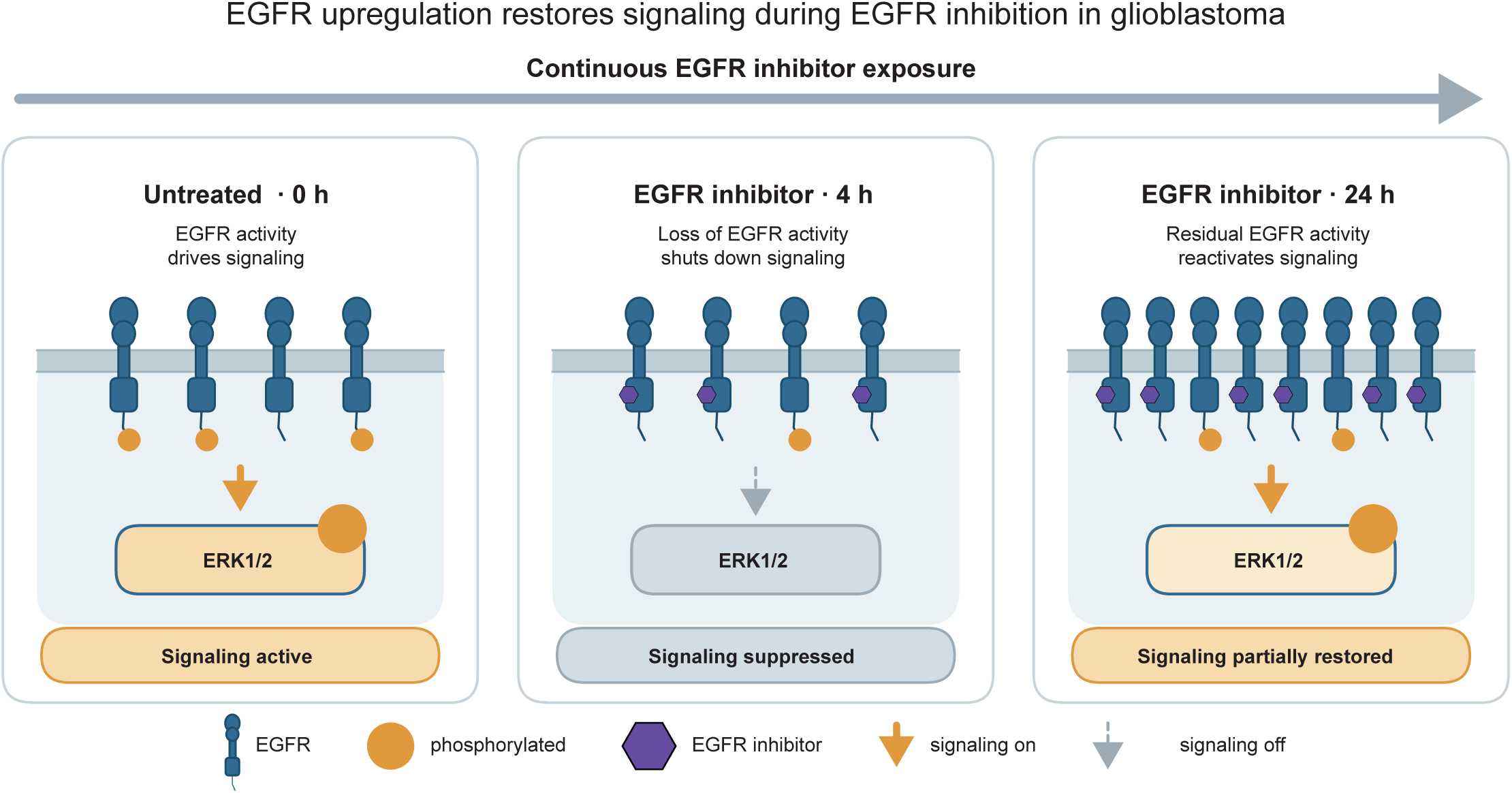

## Introduction

IDH-wildtype glioblastoma (GBM) is the most common and aggressive primary malignant brain tumor in adults^1^, with a median survival of approximately 15 months despite multimodal therapy^2–4^. Notwithstanding extensive efforts, therapeutic advances have been limited over the past two decades^5,6^. Amplification of the epidermal growth factor receptor (EGFR) gene occurs in about 40-60% of tumors and is frequently accompanied by expression of the constitutively active EGFRvIII variant^7,8^. Aberrant EGFR signaling overactivates downstream signaling pathways, such as RAS/MAPK and PI3K/AKT/mTOR, promoting cell proliferation, survival and migration^9–11^. Additional alterations affecting these pathways, including loss of chromosome 10, harboring the tumor suppressor gene phosphatase and tensin homolog (PTEN), or mutations in neurofibromin 1 (NF1), further enhance signaling output and contribute to GBM progression^12^. Aberrant EGFR signaling therefore represents a rational therapeutic target. However, EGFR-targeted therapies have shown limited clinical efficacy, despite promising preclinical activity and the availability of next-generation brain-penetrant kinase inhibitors^13–15^. In contrast to other cancers such as non-small cell lung cancer, resistance to EGFR inhibitors in GBM is not commonly driven by secondary gate-keeping mutations, suggesting that alternative mechanisms underlie therapeutic failure^10,11^. Proposed explanations include intertumoral heterogeneity, incomplete pathway inhibition, limited drug penetration, and activation of compensatory signaling pathways^16–18^.

A prevailing model is that inhibition of EGFR induces adaptive activation of compensatory signaling routes, such as expression and/or activation of alternative receptor tyrosine kinases or downstream signaling nodes, potentially enabling maintenance of pathway output through “kinome rewiring” ^19–21^. Consistent with this concept, previous studies have demonstrated signaling redundancy following targeted inhibition in GBM models^22^. However, the extent to which EGFR inhibition induces broad, kinome-wide adaptive responses remains poorly understood in GBM^23,24^. Phosphoproteomic analysis provides a systems-level approach to evaluate signaling dynamics and infer kinase activity, thus enabling the unbiased assessment of pathway adaptation following targeted inhibition^25–27^. In this study, we used phosphoproteomic profiling to characterize early signaling responses to EGFR inhibition in patient-derived GBM models. Specifically, we investigated whether EGFR inhibition induced kinome-wide adaptive responses and found instead, that the response was dominated by increased EGFR abundance that sustained residual pathway activity despite continued target suppression.

## Materials and methods

### Cell culture

Glioblastoma spheroid cultures (GSC11, GSC2, and GSC38) were kindly provided by Dr. Bhat (The University of Texas MD Anderson Cancer Center, Houston, TX, USA) and Dr. Sulman (NYU Langone’s Perlmutter Cancer Center/ NYU Grossman School of Medicine, New York, NY, USA). GBM8 was kindly provided by Dr. Bakhos Tannous (Harvard/MGH, Boston, MA, USA). BS153 was kindly provided by Dr. Zippelius (Beatrice Dolder Universitatsspital, Basel, Switzerland). VU cell lines were derived from surgical resections, with informed consent from patients, at the Amsterdam university medical centers (AUMC), location VUmc, department of neurosurgery, Amsterdam, NL), as described previously ^28^. Spheroid cultures were cultured in Neurobasal-A medium (Gibco, 10888022), supplemented with B27 minus vitamin-A (Gibco, 12587010), N-2 (Gibco, 17502001), GlutaMAX (Gibco, 35050061), EGF (20 ng/ml, Peprotech, 100-18B-1MG), bFGF (20 ng/ml, Peprotech, AF-100-15-1MG), Heparin (5000 IE/ml, VUmc pharmacy), and Penicillin/Streptomycin (Corning, 30-003-CI). BS153 was cultured in DMEM (Gibco, 11584486) with 10% Fetal bovine serum (10% v/v, VWR, S1400-500) and Penicillin/Streptomycin. All cells were cultured at 37 °C and 5% CO2 in a humidified incubator.

### DNA methylation array and copy number analysis

Genomic DNA was isolated from untreated cell pellets using the DNeasy Blood & Tissue Kit (Qiagen, 69504), according to manufacturer’s protocol. Additionally, samples were treated with RNase A (DNase and protease-free, 10 mg/mL, Thermo Scientific™, EN0531) to remove RNA from samples. EPIC methylation arrays were performed by the genomics core facility at Erasmus Medical Center, Rotterdam, NL. CNV analysis and graphing was done using the Bioconductor minfi and conumee R packages ^29^. Gene locations for annotation were derived with the UCSC genome browser database, using the Table browser tool ^30^. Assembly hg19 was used as reference genome.

### Fluorescence in situ hybridization (FISH)

Cells from GBM spheroid cultures were harvested and washed in phosphate buffered saline (PBS) pH 7.2, 0.5% bovine serum albumin (BSA) and 2 mM EDTA and diluted to 1 x 10^6^ cells/ml, as single cell suspension. 1.5 x 10^4^ cells were loaded into a cytospin cassette containing a glass microscopy slide and filter. Following centrifugation for 5 min at 150xg, cells on glass slides were air dried overnight and frozen. Fluorescence in situ hybridization was done using the ZytoLight FISH cytology implementation kit (Zytovision, Z-2099-20) with ZytoLight SPEC EGFR/CEN 7 Dual Color Probe (Zytovision, Z-2033-200), according to manufacturer’s protocol.

### Drug sensitivity assay

Cell seeding density was determined experimentally, such that growth remained in log-linear phase throughout the assay period. Drug response was determined in a 3-day drug exposure cell viability assay. Cells were seeded in 50 µL complete medium per well in 384-well plates (Greiner, 781091) and incubated according to standard culturing conditions overnight. The next day, drugs were added using a Tecan D300e automatic drug dispenser in triplicate. DMSO was normalized to 1% of the total volume in all wells except for concentrations above ∼100 µM which exceeded 1% DMSO. Cell viability was measured using CellTiter-Glo 3D (Promega, G9683), according to manufacturer’s protocol, after 72 hours of drug exposure. Response was normalized to untreated (DMSO) controls. All experiments were performed in in two independent biological replicates.

### Cell lysates preparation and immunoblotting

Cells were cultured as described and exposed to indicated drugs, or equal volume of DMSO for the indicated time and harvested for western blot analysis. After drug exposure time, cells were lysed using RIPA buffer (20mM Tris-HCL pH 7.4, 150 mM NaCl, 1mM EDTA, 1% Triton-X100, 1% sodium deoxycholate, 0.1% Sodium dodecyl sulfate (SDS) supplemented with 1 mM sodium orthovanadate and 1mM sodium fluoride), followed by vortexing and breaking up the lysate. Next, samples were centrifuged at 5400 x g for 15 min to obtain total protein extractions. Protein concentrations were measured using the Pierce BCA protein assay Kit (Thermo Fisher; #23255) according to the manufacturer’s protocol. Samples were diluted and reduced with RIPA buffer and NuPAGE™ LDS sample buffer (Thermo Fisher, #NP0007) and heated for 5 minutes at 95 °C. Equal amounts of protein, up to 100 μg per sample, and NuPAGE pre-stained protein ladder were separated via gel electrophoresis in a 4-12% Bis-Tris gradient NuPAGE gel (Thermo Fisher, #12020166) in 1x NuPAGE MOPS SDS running buffer (20x Thermo Fisher, #NP0001). Transferring of proteins onto a polyvinylidene difluoride (PVDF) membrane (Immobilon-P, Merck Millipore, Darmstadt, Germany; #IPVH00010) was performed via wet electroblotting. Antibodies used were: rabbit anti-p-Y1068-EGFR (1:1000, Cell signaling, 3777), rabbit anti-EGFR (1:1000, Cell signaling, 4267), rabbit anti-GAPDH (1:1000, Cell signaling, 2118T) and Goat anti-Rabbit–HRP secondary antibody (1:2000, Dako, P0448).

### Sample preparation LC-MS/MS

Cells were seeded at a cell density of 0.8-1.2 x 10^6^ cells/ml and cultured as described. Per condition 0.4-0.6 x 10^9^ cells were harvested to obtain ∼2.5 – 5 mg protein. Protein extraction was performed with a urea-based lysis buffer (20 mM HEPES pH 8.0, 9 M urea supplemented with 1 mM orthovanadate, 2.5 mM pyrophosphate, 1 mM β-glycerophosphate). Sample quality was assessed by global tyrosine phosphorylation levels via immunoblot with rabbit anti-p-Tyr-1000 (1:2,000, Cell signaling technology, Beverly, MA, USA) and Coomassie staining (**Figure S4**). We assessed global protein expression and phosphoprotein expression via nano-LC/MS as previously described ^27^. In short, samples were digested using trypsin and desalted with Oasis HLB 1 cm3 Vac Cartridge (Waters, 186000383). To assess phosphoprotein expression, two methods were used to enrich the phosphopeptides in the same samples prior to nano-LC/MS. First, selective capture of p-Tyr residues via immunoprecipitation with PTMScan p-Tyr-1000 Kit (Cell Signaling, 8803) was performed. Second, global capture of phosphopeptides with p-Ser, p-Thr or p-Tyr residues was performed on the AssayMAP Bravo Platform (Agilent Technologies) using 5 µl Fe(III)-NTA immobilized metal affinity chromatography (IMAC) cartridges (Agilent Technologies, G5496-60085). 200 μg desalted non-bound peptides from IP p-Tyr in 0.1% trifluoroacetic acid and 80% acetonitrile was used for this second method. Peptides were eluted in 25 µl 5% NH_4_OH/30% acetonitrile.

### LC-MS/MS

Peptides were separated using the Ultimate 3000 nano-LC–mass spectrometry (MS)/MS system (Thermo Fisher Scientific) equipped with a 50 cm × 75 μm ID Acclaim Pepmap (C18, 1.9 μm). After separation, ionization of the eluting peptides with 2 kV was performed in a Q Exactive HF mass spectrometer (Thermo Fisher Scientific) operated by Tune (v.2.11) and Xcalibur Software (v.4.3.73.11, OPTON-30965, both Thermo Fisher Scientific).

### (Phospho)-peptide quantification and data analysis

For (phospho)protein expression experiments, MS/MS spectra were referenced against the human reference proteome Fasta file (42,383 entries, canonical and isoforms, release 2021_01) using MaxQuant (v1.6.10.43) software ^31^. Trypsin-digested peptides were identified allowing up to two missed cleavage sites. Carbamidomethylation of cysteine was set as a fixed modification, while phosphorylation of serine, threonine, and tyrosine, methionine oxidation, and N-terminal acetylation were included as variable modifications. Peptide precursor ions were identified allowing a deviation of the mass up to 4.5 ppm and for fragment ions, this was 20 ppm. Peptide, protein, and site identifications were excluded based on a false discovery rate (FDR) of 1% using a decoy database strategy. The minimal peptide length for inclusion was set to seven amino acids and the Andromeda peptide modification score was set to 40, with a corresponding minimum delta score of 6 (following the default MaxQuant settings). Peptide identification was propagated across the samples using the “match between runs” option checked. Furthermore, the lysates were searched using the “label-free quantification” option selected and phospho-sites were quantified by their extracted ion intensities (“Intensity” in MaxQuant). For each sample, the phosphosite intensities were normalized (“normalized intensity”) on the median site intensity of all peptides in the sample. For global protein expression normalization was performed to the total count of each sample. Kinase activity was determined by the Integrative Inferred Kinase Activity (INKA) analysis, established by Beekhof et al, (2019) ^27^.

## Results

### Patient-derived glioblastoma models capture *EGFR*-driven molecular heterogeneity

To establish a representative model panel for downstream analyses, we first assessed whether our patient-derived GBM models recapitulates the EGFR copy number alterations observed in patient samples. Primary patient-derived cell lines were profiled using EPIC methylation arrays, enabling inference of copy number variation across a total of 29 models. EGFR amplification, defined as a chromosome 7-normalized EGFR log_2_ copy number ratio > 0.6^32^, was identified in 10 out of 29 (34.5%) cell lines (**Figure 1A-C**). To validate these findings, EGFR copy number was independently assessed by fluorescence in situ hybridization (FISH) in 16 cell lines. Using an *EGFR* to chromosome 7 ratio of more than 2 as the criterion for amplification^33^, 7 out of 15 (46.7%) cell lines were classified as *EGFR*-amplified (**Figure 1D-F**). EGFR copy number estimates obtained by FISH and array-based measurements showed a moderate correlation (r = 0.49, p = 0.033), likely reflecting methodological differences between single-cell and bulk genomic profiling. Furthermore, a number of specimens showed extrachromosomal DNA amplifications, that might show biological variation over time since their replication is independent of chromosomal replication.

**Figure 1:**
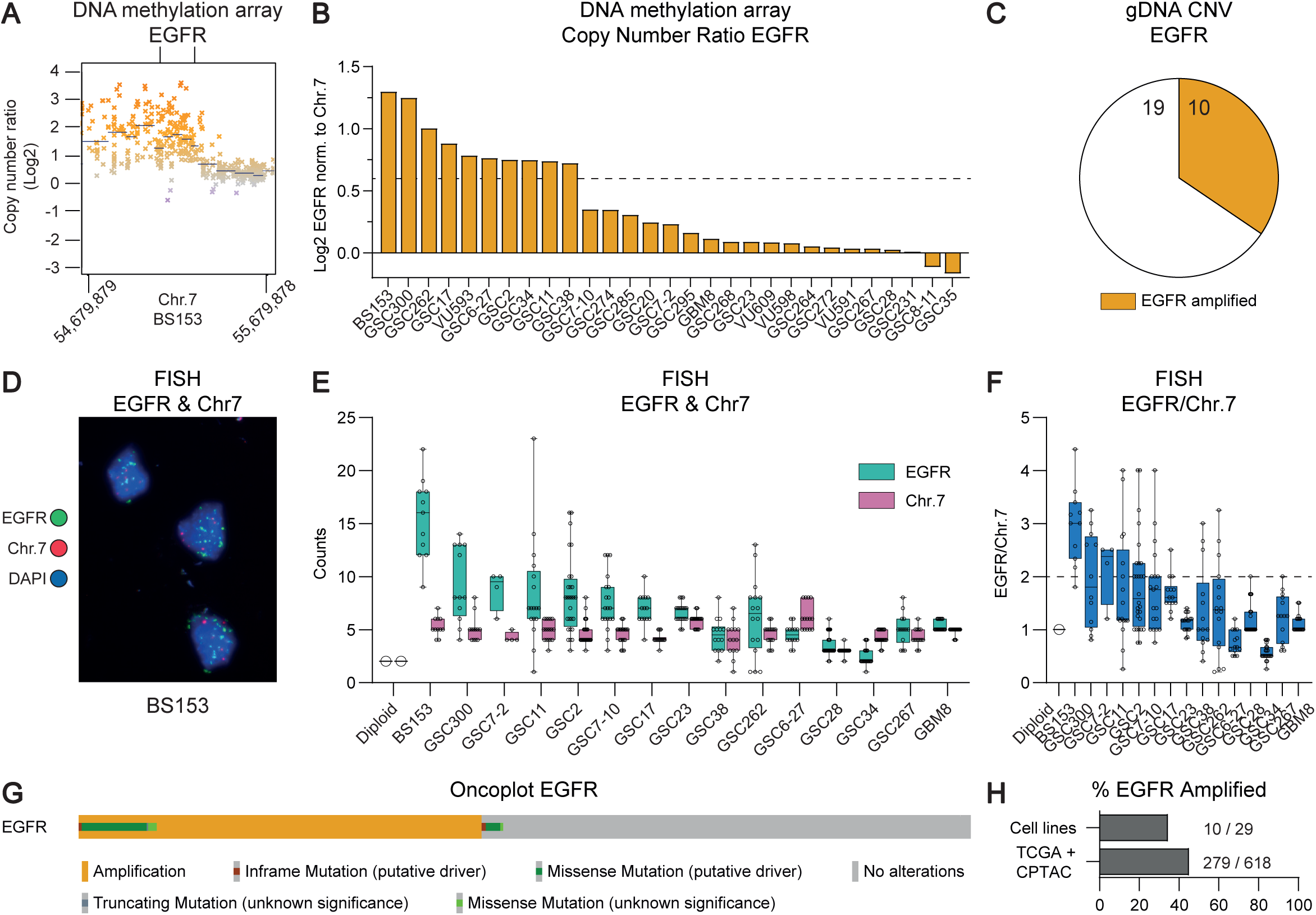
Molecular characterization of patient-derived GBM cell lines. **A)** Representative copy-number variation (CNV) profile of the *EGFR* locus in BS153 derived from EPIC DNA methylation array data showing focal *EGFR* amplification. **B)** EGFR copy-number ratios across 29 GBM cell lines, normalized to chromosome 7. A normalized log_2_ copy-number ratio > 0.6 was considered amplified. **C)** Summary of *EGFR* amplification status based on EPIC DNA methylation array-derived CNV analysis. 10 of 29 cell lines were classified as *EGFR*-amplified. Bars show average signal over the entire *EGFR* locus. **D)** Representative fluorescence in situ hybridization (FISH) image of BS153 showing *EGFR* amplification. **E)** FISH-based *EGFR* and chromosome 7 centromere counts across 15 GBM cell lines. **F)** Ratio of *EGFR* to chromosome 7 centromere signals determined by FISH. An *EGFR*/Chr.7 ratio > 2 was considered amplified. In E) and F), boxes indicate the interquartile range with the median shown as the center line, and dots represent individual measurements. **G)** Oncoplot showing *EGFR* alterations found in GBM tumors from TCGA and CPTAC. Data were obtained from cBioPortal. **H)** Frequency of *EGFR* amplification in the patient-derived GBM cell line panel compared with TCGA/CPTAC GBM tumors.

Consistent with clinical GBM cohorts, chromosome 7 gain frequently co-occurred with increased EGFR copy number in the majority of cell lines (**Figure 1D-F**). Overall, the frequency and distribution of EGFR alterations in this model panel closely mirrors those reported in large GBM patient cohorts, including The Cancer Genome Atlas (TCGA)^11^ and Clinical Proteomic Tumor Analysis Consortium (CPTAC)^10^ (**Figure 1G,H**).

Based on these genomic analyses, five representative cell lines spanning *EGFR* amplification states were selected for further analysis, including two with high *EGFR* amplification (BS153, GSC11), two with moderate *EGFR* amplification (GSC38, GSC2), and one lacking detectable *EGFR* aberration (GBM8). Cell lines with high *EGFR* amplification displayed greater intercellular heterogeneity in both EPIC array and FISH analyses, while moderately *EGFR*-amplified, and *EGFR*-low models showed more homogenous copy number profiles (**Figure 1; Figure S1**).

### Distinct basal signaling states characterize patient-derived GBM models

To further characterize the selected models, we performed global proteomic and phosphoproteomic profiling using LC-MS/MS, enabling quantification of protein expression and inference of kinase activity. A principal component analysis (PCA) of kinase activity profiles, obtained using Integrative Inferred Kinase Activity (INKA)^27^ analysis, revealed that cell lines grouped according to their overall signaling state rather than EGFR activity alone (**Figure 2A**). Indeed, inspection of the components indicated that, contrary to our expectations, neither PC1 nor PC2 was primarily driven by EGFR signaling. Nevertheless, EGFR remained one of the most dominant signaling nodes in cell lines with high *EGFR* amplification (**Figure S2**). Other GBM-relevant receptor tyrosine kinases, including PDGFRA, a hallmark of the proneural GBM subtype^34^, INSR, which promotes glioblastoma cell proliferation and survival^35^, and FGFR1, which contributes to glioblastoma invasion^36^, also contributed to the signaling landscape across the model panel. Pathway mapping based on INKA scores, with network visualization of kinase-substrate components, further showed comparable EGFR signaling activity among cell lines with comparable *EGFR* amplification status (**Figure 2B**). In contrast, EGFR activity was not detected in GBM8, in which PDGFRA represented the predominant signaling node, supporting its use as an EGFR-independent reference model (**Figure 2B**).

**Figure 2:**
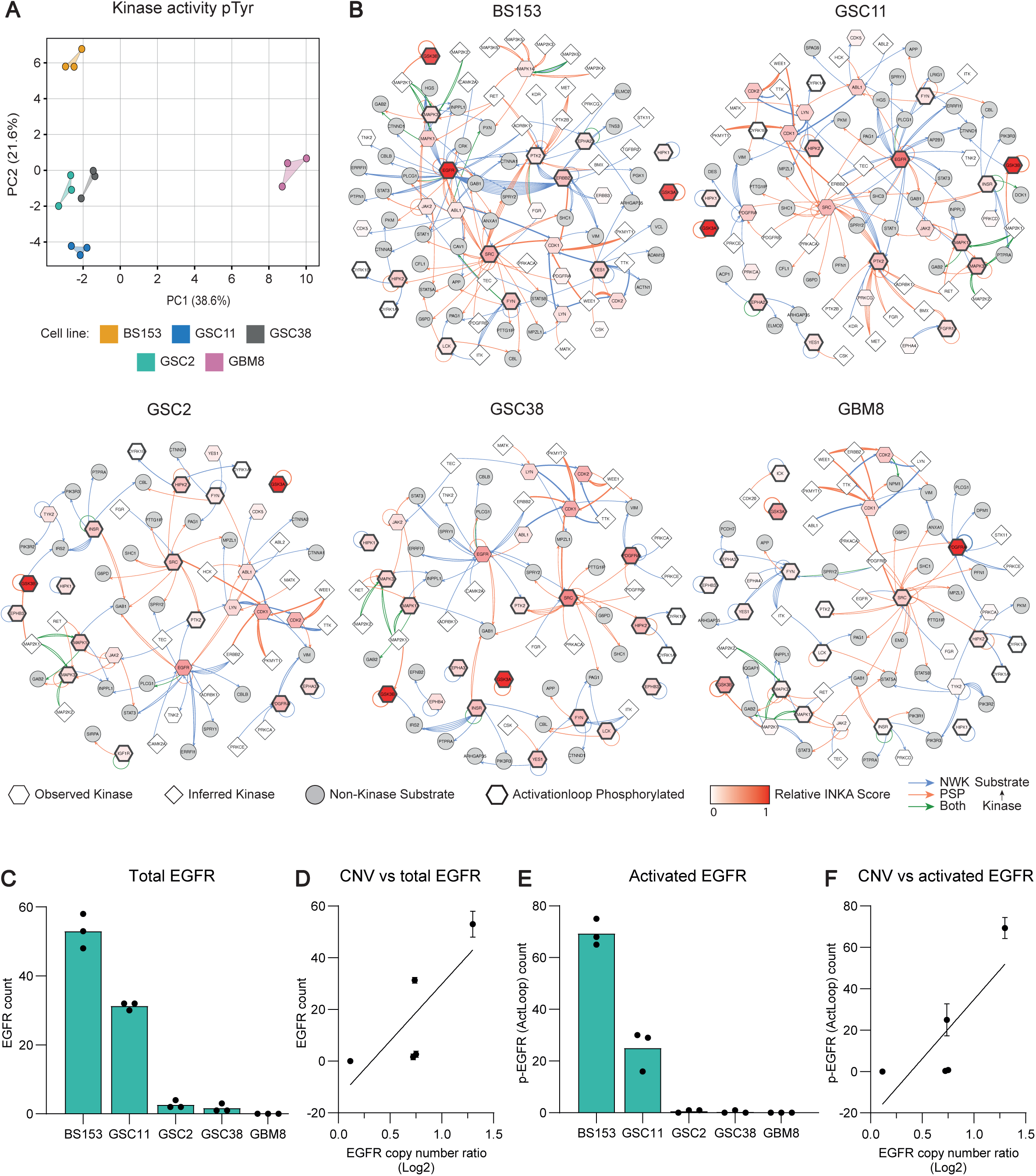
Basal signaling states and EGFR activity in patient-derived GBM models. **A)** Principal component analysis (PCA) of basal kinase activity inferred from phosphotyrosine (pTyr) phosphoproteomic data using INKA. Colors indicate individual GBM cell lines (BS153, orange; GSC11, blue; GSC38, gray; GSC2, teal; GBM8, magenta). **B)** INKA-derived signaling networks for the five selected GBM cell lines. Nodes represent observed kinases (hexagon), inferred kinases (diamond), and non-kinase substrates (circle); a thick outline indicates activation loop phosphorylated kinases. Node color indicates relative INKA score, and kinase-substrate relationship are represented by arrows (blue, relationship predicted by NetworKIN (NWK); red, relationship experimentally observed provided by PhosphoSitePlus (PSP); green, both). **C)** Total EGFR abundance measured by LC-MS/MS across the five cell lines. **D)** Correlation between total EGFR abundance and *EGFR* copy number ratio (Spearman r = 0.9). **E)** EGFR phosphorylation at activation loop (Act Loop) sites measured by LC-MS/MS across the five GBM cell lines. **F)** Correlation between activation loop EGFR phosphorylation and *EGFR* copy number ratio (Spearman r = 0.9). In C) and E), bars represent the mean, and dots indicate individual data points while in D) and F), data represent the mean +/-SD (N=3).

### *EGFR* gene amplification is associated with protein abundance and activity

Given the prominent role of EGFR signaling in *EGFR*-amplified GBM models, we next examined whether *EGFR* gene amplification was associated with receptor abundance and activation. For this purpose, total EGFR protein levels and phosphorylation were quantified using LC-MS/MS (**Figure 2C**). This analysis revealed that EGFR protein abundance increased with higher *EGFR* gene copy number across the cell line panel, showing a strong positive correlation between gene amplification and protein expression (Spearman r = 0.9; **Figure 2D**). These findings indicate that increased *EGFR* gene copy number is associated with higher receptor abundance. Furthermore, EGFR phosphorylation on residues in the activation loop also scaled with EGFR protein abundance or copy number (Spearman r = 0.9; **Figure 2E-F**).

### EGFR signaling partially recovers during sustained EGFR inhibition

To investigate the cellular response to EGFR inhibition, the selected GBM models were analyzed using viability measurements, immunoblotting, and phosphoproteomic profiling. In order to identify a suitable drug concentration for reducing viability while preserving sufficient cells for downstream analyses, cells were treated with increasing concentrations of the structurally different, brain penetrant EGFR inhibitors, Osimertinib and Zorifertinib. Cell viability was measured after 72 hours of drug exposure (**Figure S3**). Both inhibitors reduced cell viability of EGFR driven GBM cells by approximately 50% at concentrations of ∼0.5 µM (**Figure 3A**). Sensitivity to EGFR inhibition closely correlated with EGFR activation (**Figure 2E**), suggesting that basal receptor activity is a good predictor of drug response. These results prompted us to examine EGFR signaling dynamics over time following inhibitor exposure. Immunoblot analysis of cells treated with 0.5 µM Zorifertinib showed that EGFR phosphorylation was strongly suppressed for the first 7 hours of treatment, but re-increased progressively and partially recovered after 24 hours. This recovery coincided with an increase in total EGFR abundance (**Figure 3B**).

**Figure 3:**
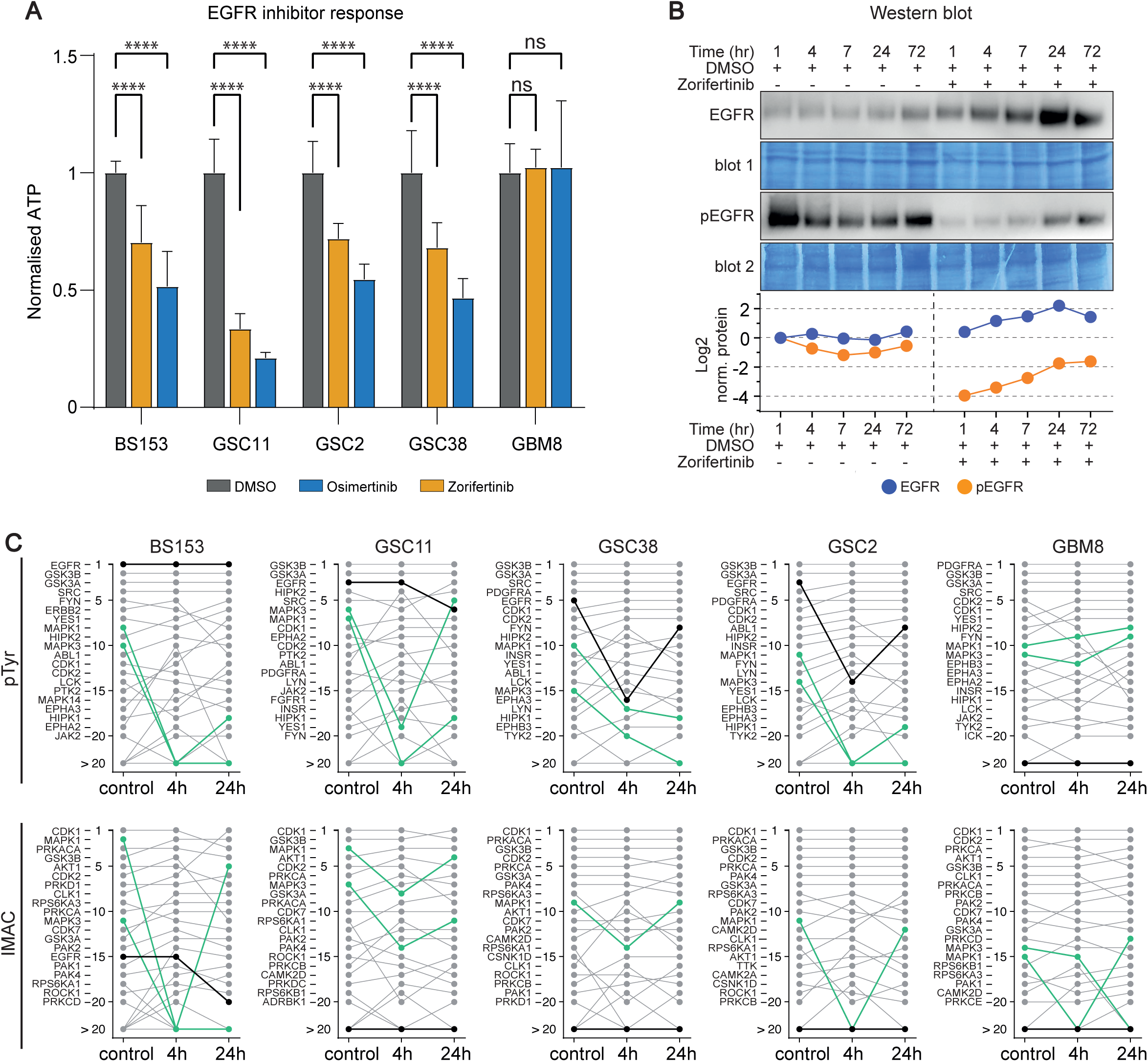
EGFR signaling partially recovers during sustained EGFR inhibition. **A)** Viability of the indicated GBM cell lines following treatment with osimertinib and zorifertinib. Cell viability was measured by ATP quantification after 72 h and normalized to DMSO-treated controls. Bar shows mean ±SD at ∼0.5 µM (Statistical significance was determined by two-way ANOVA; **** = p < 0.0001; ns, not significant (N = 6). **B)** Immunoblot analysis of total and phosphorylated EGFR (pEGFR Y1068) in GSC2 cells during treatment with 0.5 µM zorifertinib) or DMSO for up to 72 hours. Bottom panel shows log_2-_normalized EGFR and pEGFR signal relative to the 1h DMSO control. **C)** Time-resolved kinase activity inferred by INKA from pTyr-and IMAC-enriched phosphoproteomic data at baseline/control, 4 h, and 24 h after EGFR inhibition. Kinases are ranked according to their INKA score in control samples and followed across treatment. EGFR is indicated with a thick black line and MAPK1 (ERK2) and MAPK3 (ERK1) in green.

To further characterize the signaling changes accompanying re-activation of the EGFR pathway, we performed phosphoproteomic profiling at early (4 hours) and late (24 hours) time points, corresponding to maximal inhibition and partial signaling recovery, respectively. INKA-based Kinase activity analysis showed reduced activity of EGFR and its downstream effectors, including MAPK1/3 (ERK2/1), after 4 hours of treatment, consistent with effective pathway inhibition (**Figure 3C; Table S1-S2**). By 24 hours, EGFR activity and downstream signaling activity had partially recovered across multiple cell lines. Recovery of EGFR activity was most apparent in the moderately amplified models (GSC38, GSC2), whereas downstream pathway suppression and subsequent recovery was more pronounced in higher EGFR-amplified cell lines (BS153, GSC11).

Beyond EGFR and its downstream signaling components, only limited changes in kinase activity were observed, suggesting that adaptive responses were largely confined to the EGFR pathway in this context. Interestingly, although *PDGFRA* amplification is a defining feature of the proneural GBM subtype^34^ and PDGFRA-driven signaling programs are characteristic of EGFR-independent GBM^37^, it consistently ranked among the most active kinases across multiple EGFR-driven cell lines. However, EGFR inhibition did not result in compensatory activation of PDGFRA signaling in these models. These results show that early adaptation to EGFR inhibition is characterized by partial restoration of EGFR pathway signaling rather than widespread activation of alternative kinase pathways.

### EGFR inhibition does not induce broad kinome rewiring

To formally assess whether EGFR pathway reactivation was accompanied by broader adaptive changes in kinase activity, we next examined global kinome responses following EGFR inhibition. Previous studies have shown that targeted kinase inhibition can induce compensatory activation of alternative signaling pathways, including large-scale remodeling of kinase activity networks (“kinome rewiring”), as revealed by phosphoproteomic analyses in multiple cancer models and contexts of EGFR inhibition. However, whether such broad remodeling occurs during early responses to EGFR inhibition in GBM remains unclear.

Pairwise correlation analysis between inhibitor-treated and control samples showed high concordance across all conditions (**Figure 4A,B**). The largest differences were mainly confined to EGFR and its downstream effectors, consistent with on-target pathway inhibition. Similarly, PCA analysis of kinase activity profiles revealed that samples grouped primarily by cell line identity rather than treatment condition, with EGFR inhibitor-treated and untreated samples remaining closely grouped together within each model (**Figure 4C**). Comparable patterns were observed across *EGFR*-high, *EGFR-*moderate, and EGFR-independent cell lines, indicating that inhibitor treatment does not drive large-scale restructuring of the kinome activity. Consistent with these findings, PCA performed separately for each cell line did not reveal treatment-specific grouping (**Figure S5**).

**Figure 4:**
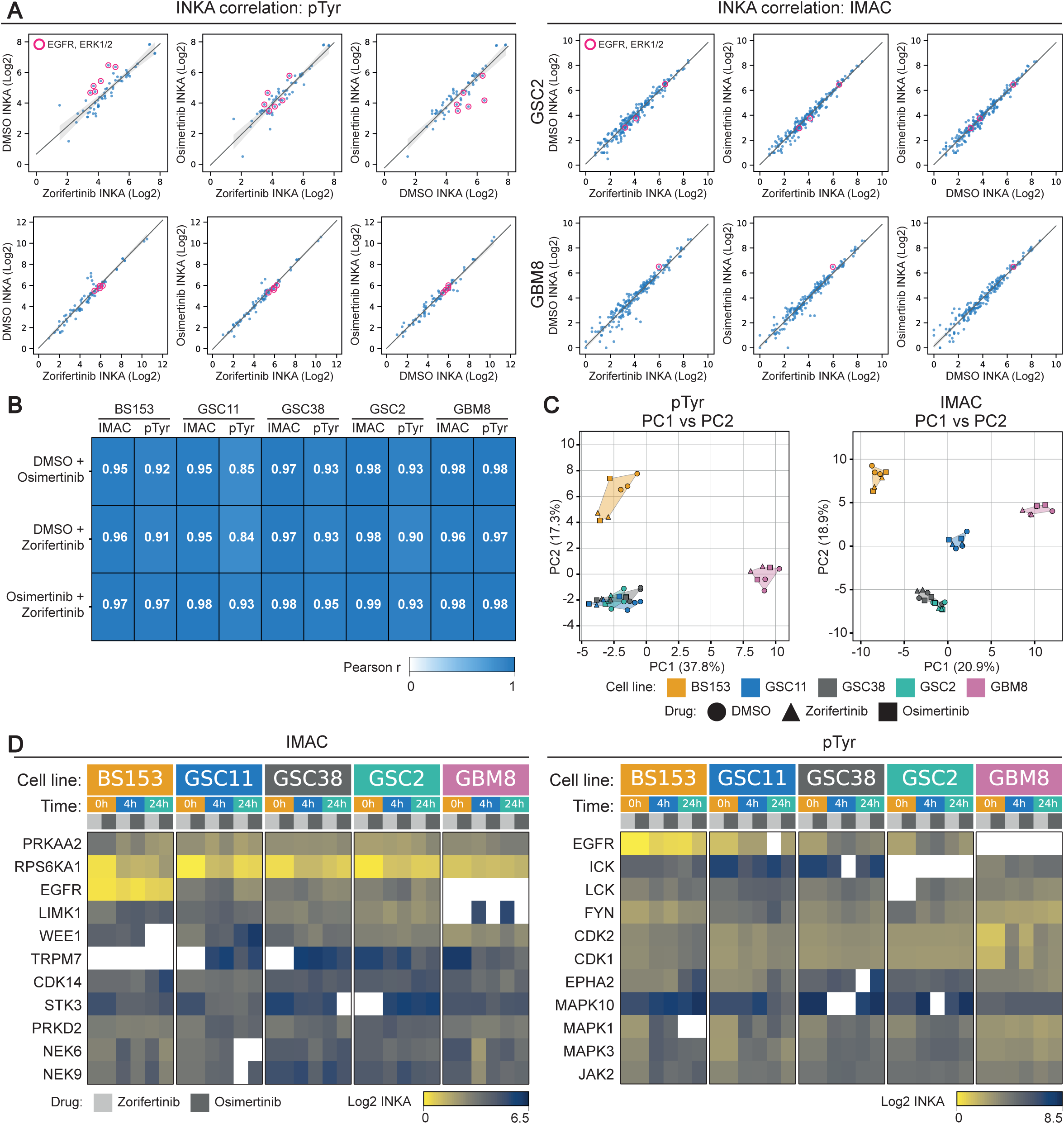
EGFR inhibition does not induce broad kinome rewiring. **A)** Pairwise comparison of log_2_-transformed INKA scores between DMSO-, osimertinib-, and zorifertinib-treated samples for pTyr-and IMAC-enriched phosphoproteomic datasets. EGFR and ERK1/2 are highlighted in magenta. **B)** Pearson correlation coefficients between INKA profiles for the indicated treatment conditions in each cell line for pTyr-and IMAC-enriched datasets. **C)** Principal component analysis of INKA-derived kinase activity profiles from pTyr-and IMAC-enriched phosphoproteomic data. Colors indicate cell lines (BS153, orange; GSC11, blue; GSC38, gray; GSC2, teal; GBM8, magenta), and symbols indicate treatment (DMSO, circle; zorifertinib, triangle; osimertinib, square). **D)** Heatmaps showing log_2_ INKA scores for kinases displaying prominent changes across treatment conditions in the IMAC-and pTyr-enriched datasets at 0 h (orange), 4 h (blue), and 24 h (green) of treatment with zorifertinib (light grey) or osimertinib (dark grey). White indicates kinase for which no INKA score was obtained.

To further examine localized changes in kinase activity, kinases were ranked according to differential INKA scores following inhibitor treatment. This analysis revealed that changes were indeed predominantly restricted to EGFR and its immediate downstream signaling components, including ERK1/2, JAK2, and S6K1 (**Figure 4D**). Outside of this pathway, only minor alterations in kinase activity were observed. As expected, the EGFR-independent cell line GBM8 showed no detectable EGFR activity and minimal changes in downstream signaling upon inhibitor treatment. These data indicate that EGFR inhibition does not induce broad kinome rewiring during the first 24 hours of treatment in these models but instead results in pathway-restricted signaling changes largely confined to the EGFR pathway itself.

### Increase of EGFR abundance is associated with partial restoration of signaling during sustained inhibition

To determine whether adaptive responses occur within the EGFR pathway and to investigate the mechanism underlying the partial signaling recovery, we examined changes in EGFR abundance and phosphorylation during sustained inhibitor exposure. Cells were treated with 0.5 μM Zorifertinib for 4 and 24 hours, after which total and phosphorylated EGFR levels were compared to DMSO-treated controls by immunoblotting and LC-MS/MS.

Consistent with the time-course analysis (**Figure 3B**), total EGFR levels increased after 24 hours of treatment (**Figure 5A,B**). This increase was accompanied by higher absolute levels in EGFR phosphorylation in all EGFR-dependent cell lines tested. To determine whether receptor activation changed relative to protein abundance, the ratio of phosphorylated to total EGFR was quantified from immunoblot analyses. This ratio remained largely unchanged between 4 and 24 hours of treatment (p > 0.05), indicating that the proportion of phosphorylated receptor remained constant despite increased EGFR abundance. Together, these findings suggest accumulation of EGFR protein is associated with the partial restoration of downstream signaling observed during sustained inhibitor treatment, while remaining consistent with continued target engagement by the inhibitor.

**Figure 5:**
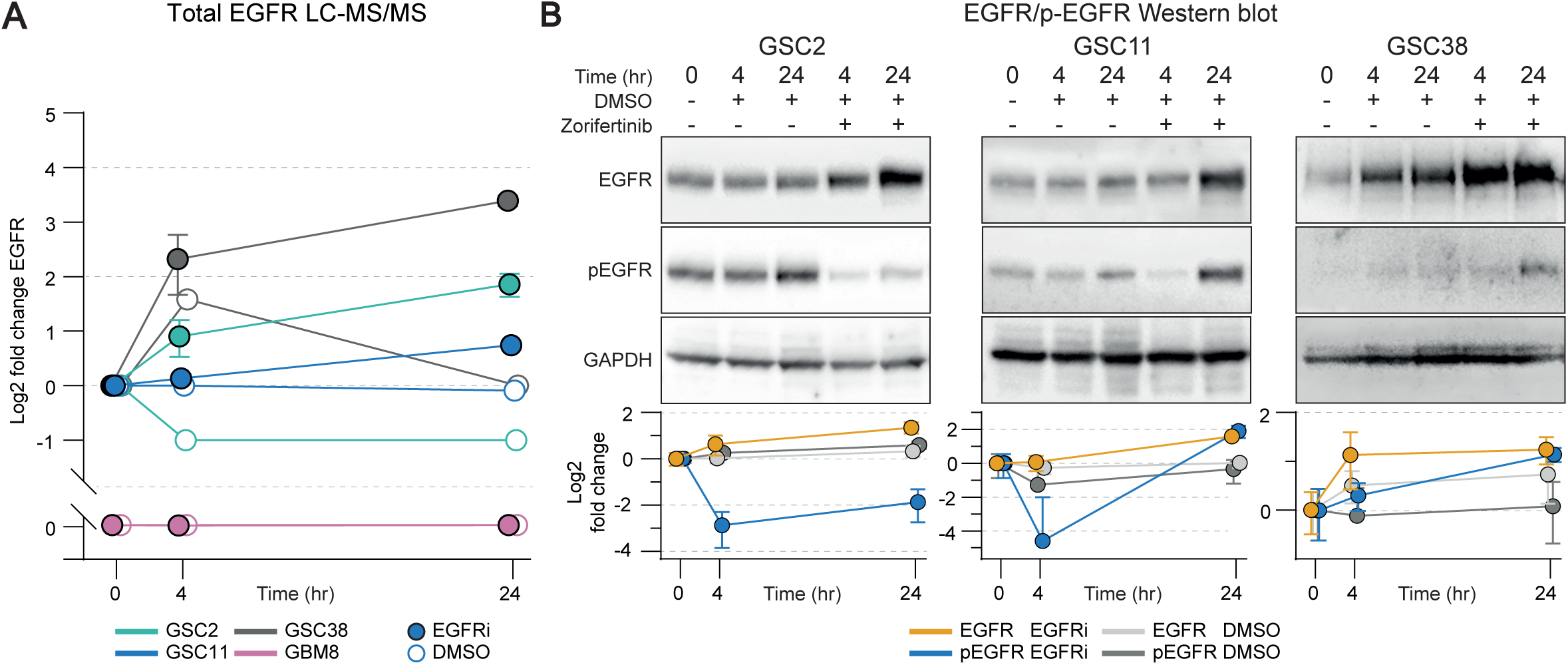
EGFR abundance and residual phosphorylation increase during sustained EGFR inhibition. **A)** Changes in total EGFR abundance measured by LC-MS/MS after 4 and 24 h of EGFR inhibitor treatment. Values are shown as log_2_ fold change relative to untreated controls for the indicated cell lines. Data are presented as the geometric mean ±SD. **B)** Immunoblot analysis of total EGFR and phosphorylated EGFR (pEGFR Y1068) in GSC2, GSC38, and GSC11 cells following 4 and 24 h of zorifertinib treatment, with corresponding DMSO-treated controls. GAPDH was used as a loading control. Graphs show log_2_ fold changes in total EGFR and pEGFR relative to baseline. Data are presented as geometric mean ± SD (N = 4).

## Discussion

In this study, we combined genomic, proteomic and phosphoproteomic analyses of patient-derived glioblastoma models to investigate early adaptive responses to EGFR inhibition. We found that early adaptation was characterized by partial restoration of EGFR signaling without extensive remodeling of global kinase activity. Furthermore, increased EGFR protein abundance was associated with this restoration of signaling despite continued inhibitor exposure.

EGFR amplification is one of the most frequent genomic alterations in glioblastoma and therefore a major therapeutic target. Consistent with its clinical relevance, our patient-derived models recapitulated the spectrum of EGFR amplification observed in patient populations, providing a representative system to investigate responses to EGFR-targeted therapy. Despite decades of effort, however, EGFR inhibitors have shown limited clinical benefit in glioblastoma.

Early-generation EGFR inhibitors were limited by poor brain penetration, whereas newer compounds such as Osimertinib and Zorifertinib-developed for lung cancers harboring gate-keeping mutations like T790M while also targeting brain metastases - achieved therapeutically relevant concentrations in the central nervous system^13–15^. Despite improved brain penetration, responses to EGFR inhibition in glioblastoma remain limited, suggesting that intrinsic adaptive mechanisms contribute to therapeutic resistance. One proposed mechanism is activation of compensatory kinase signaling, often referred to as kinome rewiring. However, our phosphoproteomic analyses provide little evidence for widespread remodeling of kinase activity, at least during the first 24 hours of EGFR inhibition.

Adaptive responses to kinase inhibition, often referred to as kinome rewiring, have been reported in several cancer models upon specific kinase inhibitor treatment. For example, Lin et al. described adaptive changes in mRNA expression, protein abundance, and inhibitor-binding profiles, and identified increased sensitivity to combined EGFR and CDK6 inhibition^38^. While these findings demonstrate substantial molecular adaptation, they did not directly assess kinase activity. In contrast, our phosphoproteomic analyses revealed only limited changes in kinase activity following EGFR inhibition during the first 24 hours of EGFR inhibition, with signaling alterations remaining largely confined to the EGFR pathway. These findings suggest that early proteomic and/or transcriptional adaptations may not necessarily coincide with widespread functional remodeling of kinase signaling output.

Although several receptor tyrosine kinases, like PDGFRA, FGFR1, INSR and others were active in our EGFR-dependent GBM models, we found no evidence that these receptors became preferentially activated to compensate for the loss of EGFR signaling. This contrast with previous studies that have described involvement of other kinases (mostly receptor tyrosine kinases) in resistance to EGFR inhibitors in GBM, none have resulted in a functional therapeutic combination^39^. While such pathways may contribute to signaling under specific conditions, our phosphoproteomic analyses suggest that they do not constitute a dominant mechanism of early adaptation in the models studied here.

Alternative mechanisms of resistance to receptor tyrosine kinase inhibitors involve altered receptor trafficking and degradation. For instance, a recent study in non-small cell lung cancer showed that disruption of clathrin-mediated endocytosis rerouted mutant EGFR towards macropinocytosis and lysosomal degradation, resulting in loss of EGFR signaling and enhanced sensitivity to EGFR inhibitors^40^. These findings highlight that changes in EGFR turnover can strongly influence drug response. Although the underlying mechanisms remain to be determined, and will be subject of future studies, a similar process could contribute to the adaptive responses observed in glioblastoma.

In our study, total EGFR abundance progressively increased during sustained inhibitor exposure and was accompanied by partial restoration of downstream EGFR and ERK signaling. The ratio of phosphorylated to total EGFR remained largely unchanged between 4 and 24 hours, consistent with continued but incomplete inhibition despite increased receptor abundance. One possible explanation is that the accumulation of EGFR raises residual kinase activity above the threshold required to activate downstream signaling, allowing partial restoration of pathway activity without widespread remodeling of kinase networks. However, our data do not distinguish whether EGFR accumulation results from enhanced transcription or translation, reduced receptor degradation, altered trafficking, or a combination of these mechanisms. The increase in EGFR protein expression may reflect adaptive pathway rescaling in response to acute suppression of EGFR signaling. Similar feedback responses have been described following growth factor deprivation and restoration, where increased cell-surface receptor abundance restores signaling sensitivity^41^. Consistent with this concept, several regulators of EGFR signaling and receptor turnover, including MIG6 (ERRFI1) and SPRY proteins, have been implicated in feedback regulation of EGFR abundance and activity^42,43^. Interestingly, these proteins were associated with EGFR signaling in our INKA network analysis (**Figure 2B; Table S1-S2**). Whether they contribute to the changes in EGFR abundance observed here remains unknown, but their established roles in EGFR feedback regulation provide a mechanistic framework for future studies investigating how EGFR abundance is regulated during sustained inhibitor treatment.

In summary, our study demonstrates that early adaptation to EGFR inhibition in patient-derived GBM models occurs without widespread remodeling of kinase activity. Partial restoration of EGFR signaling is accompanied by increased EGFR protein abundance despite continued inhibitor exposure. These findings suggest that regulation of receptor abundance, rather than activation of alternative kinase pathways, may represent an important component of the early adaptive response to EGFR inhibition. Determining how EGFR abundance is regulated following inhibitor treatment and whether this process can be therapeutically targeted will be an important direction for future studies.

## Acknowledgments

Dr. Sulman (NYU Langone’s Perlmutter Cancer Center/ NYU Grossman School of Medicine, New York, NY, USA); Dr. Bakhos Tannous (Harvard/MGH, Boston, MA, USA) and Dr. Zippelius (Beatrice Dolder Universitatsspital, Basel, Switzerland) are thanked for providing relevant cell culture models.

## Required statements

### Ethics

Tumor sample collection at MD Anderson was performed under protocol #LAB03-0687, approved by the institutional review board, after written informed consent was obtained from the patients. Tumor samples at the Amsterdam UMC and UMC Groningen were collected under the National policy on further use of bodily material CCMO #7:467.

### Authorship

YB performed experiments, formal analyses, writing manuscript; MH performed experiments, formal analyses; TTW, PP, RGH and SP performed experiments; TP and AH bioinformatics analyses; CJ and DN, supervision; AG supervision, conception and writing manuscript; BAW supervision, conception and funding. All authors read the manuscript.

### Data availability

The raw DNA methylation array data are publicly available in the Gene Expression Omnibus (GEO) under accession number GSE342610.

## Supplementary figure legends

**Figure S1:**
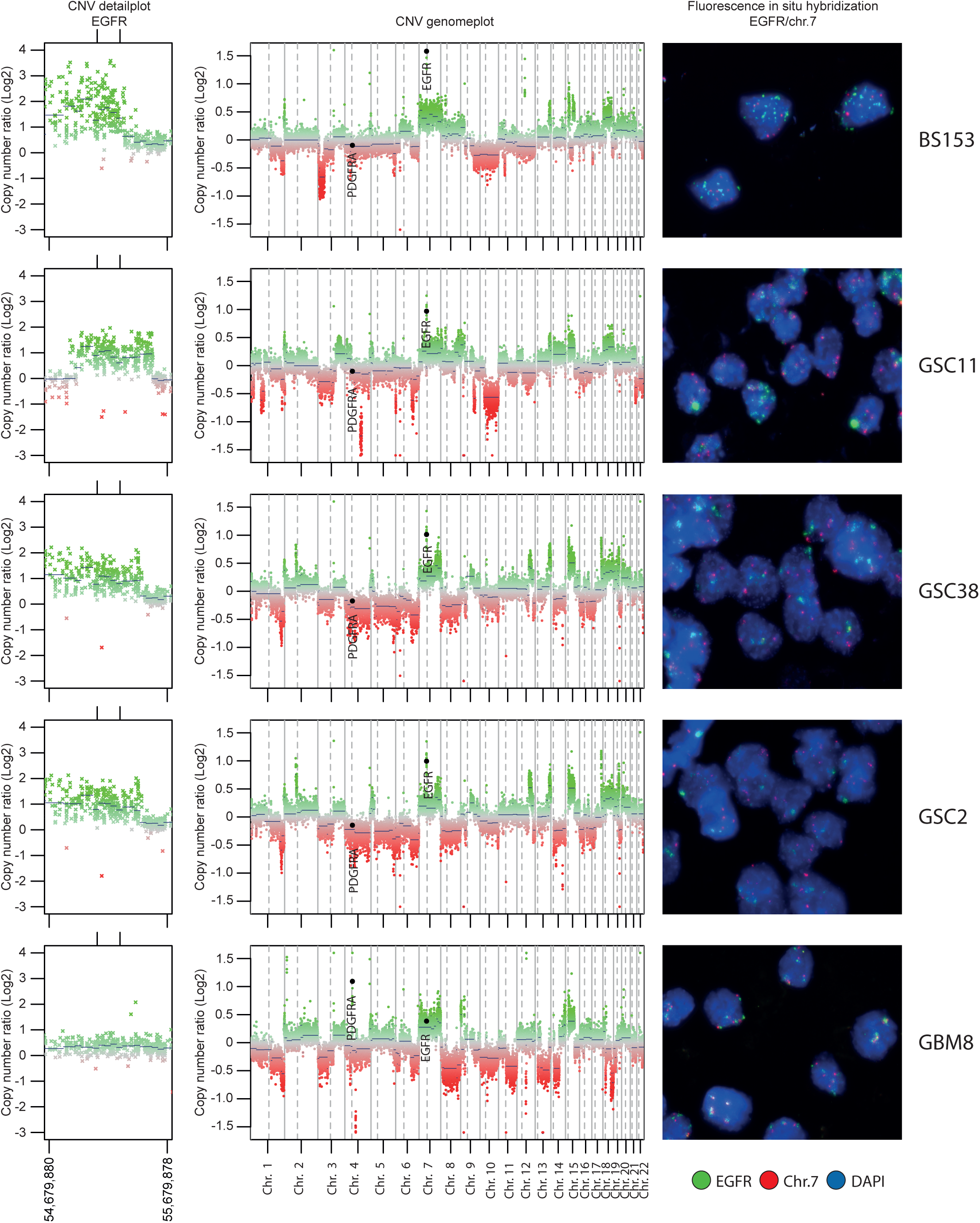
Molecular Characterization of selected GBM cell lines. CNV detail profile of the *EGFR* locus (left), genome-wide CNV profiles highlighting *EGFR* and *PDGFRA* (middle), and representative *EGFR*/Chr.7 FISH images (right) for BS153, GSC11, GSC2, GSC38, GBM8. In FISH images, *EGFR* is shown in green, chromosome 7 centromere in red, and nuclei (DAPI) in blue.

**Figure S2:**
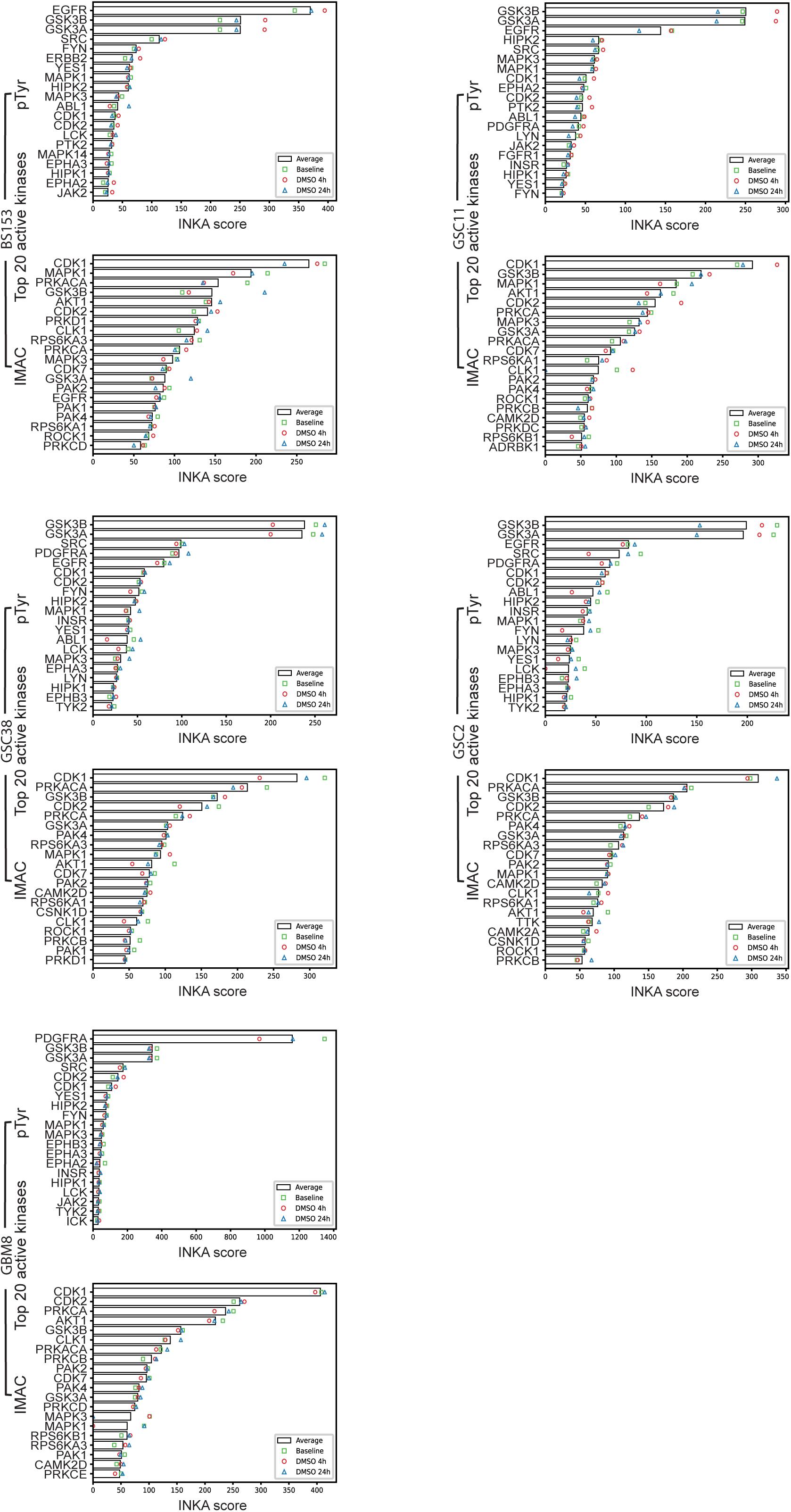
Basal kinase activity across GBM models. Top 20 active kinases ranked by INKA score in untreated BS153, GSC11, GSC38, GSC2, and GBM8 cells, derived independently from pTyr-and IMAC-enriched phosphoproteomic datasets.

**Figure S3:**
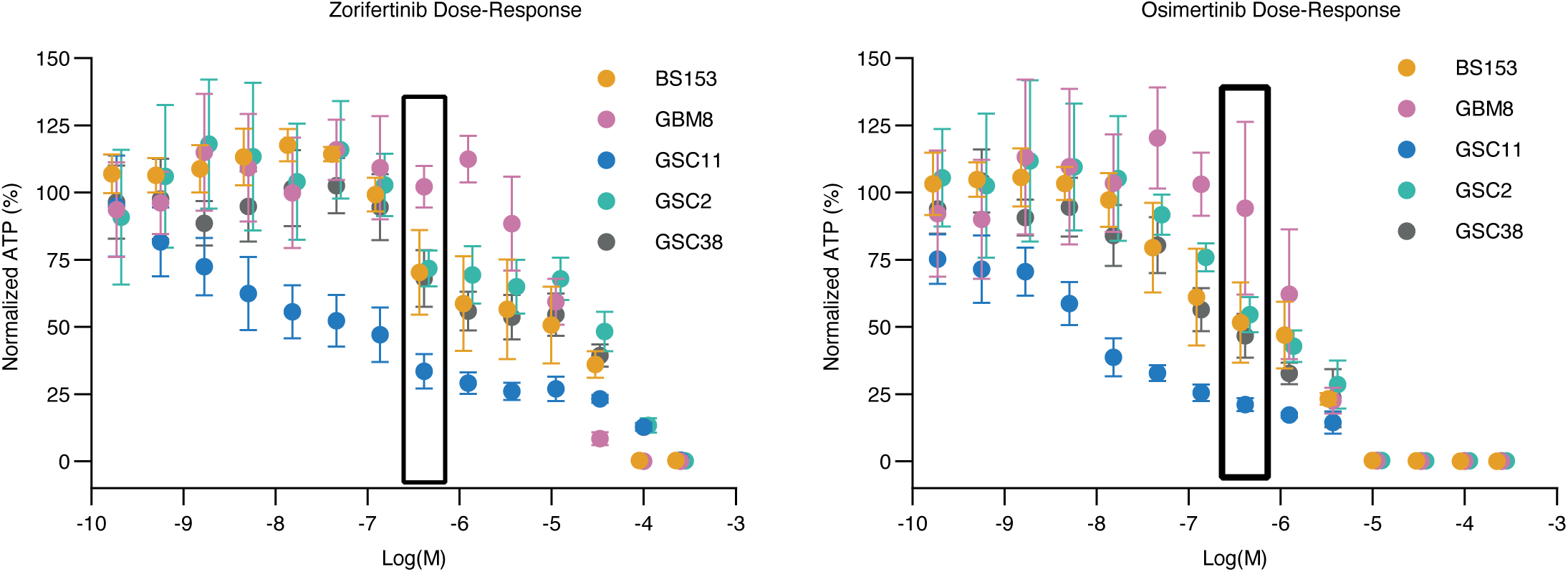
Dose-response to EGFR inhibitors. Dose-response curves for zorifertinib and osimertinib across the five selected GBM cell lines. Cell viability was measured by ATP quantification after 72 h of treatment and normalized to untreated controls. Box indicates the concentration range from which ∼0.5 µM was selected for subsequent experiments

**Figure S4:**
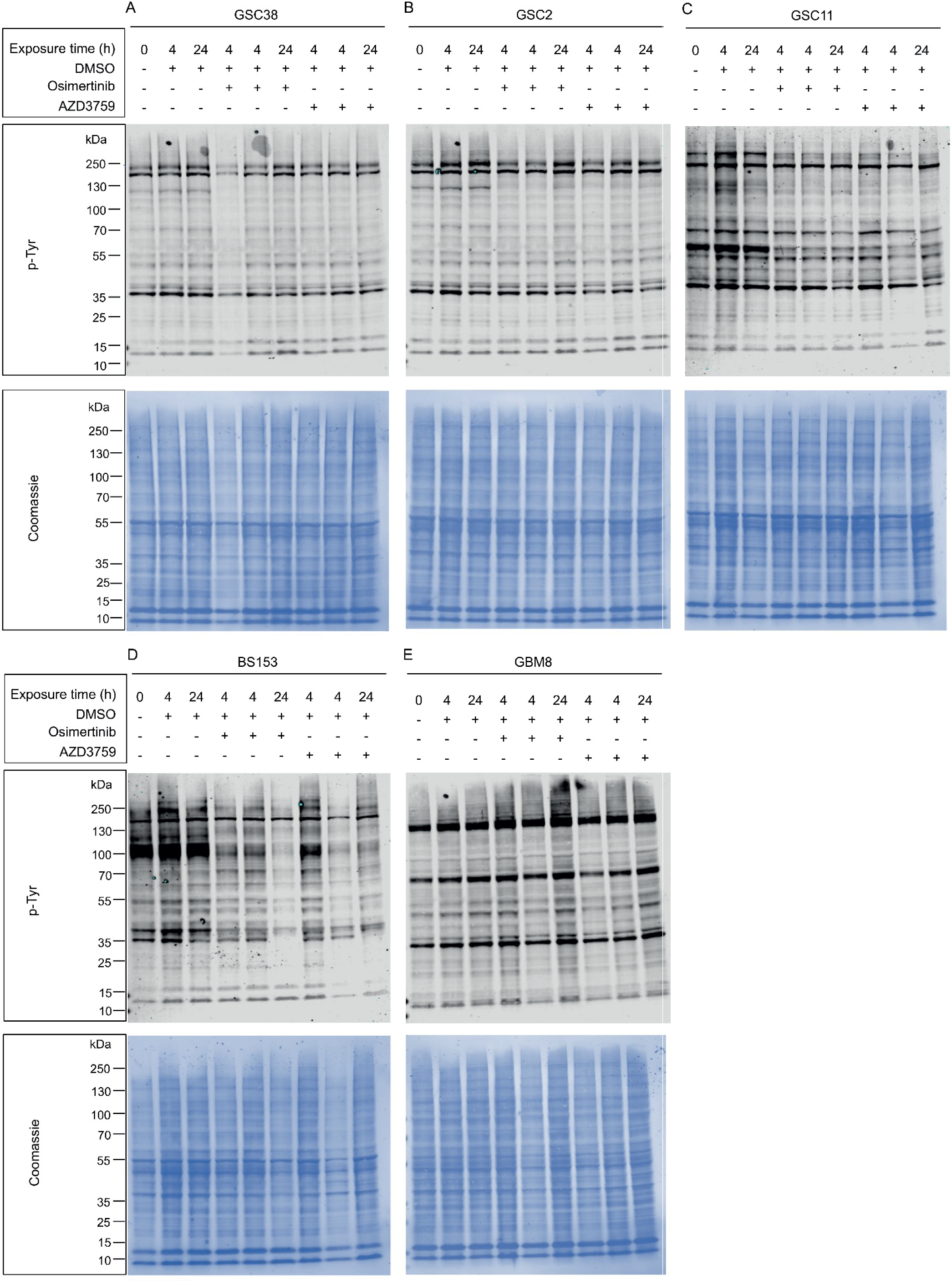
Quality control of samples used for LC-MS/MS phosphoproteomic analysis. Global tyrosine phosphorylation was assessed by anti-phosphotyrosine immunoblotting in samples collected at the indicated time points following DMSO, osimertinib, or zorifertinib treatment. Coomassie staining shows total protein loading.

**Figure S5:**
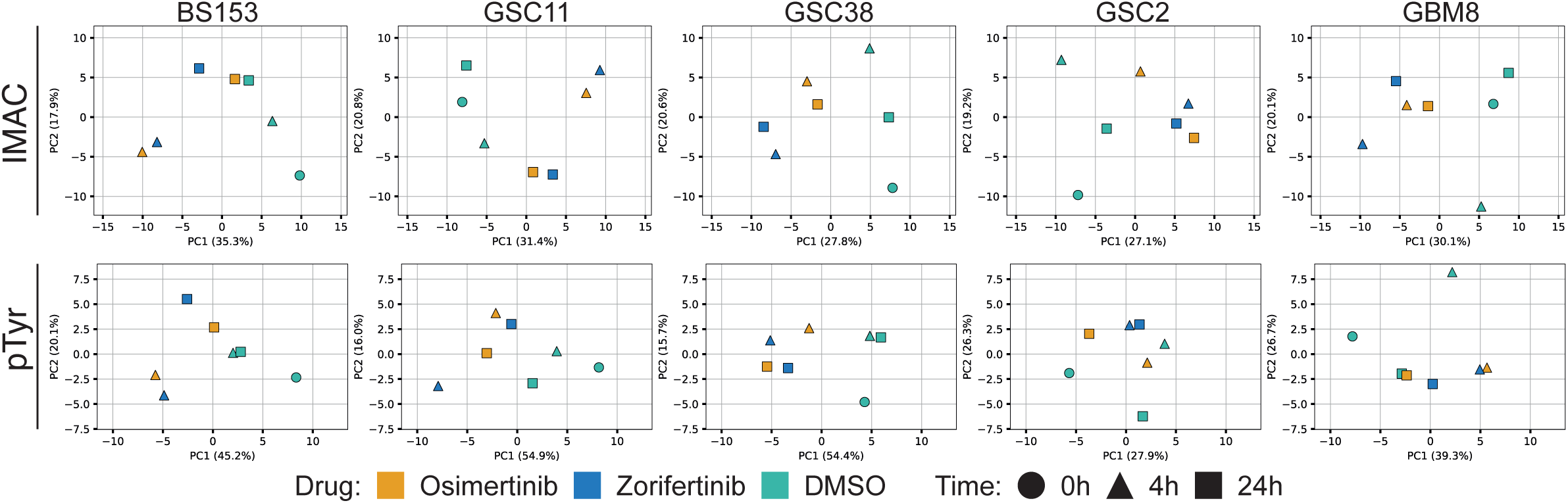
Principal component analysis of treatment responses within individual GBM cell lines. PCA of INKA-derived kinase activity profiles performed separately for each cell line using IMAC-and pTyr-enriched phosphoproteomic datasets. Colors indicate treatment and symbols indicate time point (0, 4, or 24 h).

